# Internally-controlled and dynamic optical measures of functional tumor biology

**DOI:** 10.1101/2022.05.30.493733

**Authors:** Taemoon Chung, Libia Garcia, Manojit M. Swamynathan, Fieke E.M. Froeling, Lloyd C. Trotman, David A. Tuveson, Scott K. Lyons

**Author notes:** Corresponding author: Scott K. Lyons.

## Abstract

Imaging defined aspects of functional tumor biology with bioluminescent reporter transgenes is a popular approach amongst the research community in drug development, as it is sensitive, relatively high-throughput and low cost. However, the lack of internal controls subject functional bioluminescence to a number of unpredictable variables that reduce this powerful tool to semi-quantitative interpretation of large-scale effects. Here we report the generation of sensitive and quantitative live reporters for two key measures of functional cancer biology and pharmacologic stress: the cell cycle and oxidative stress. We developed a two-colored readout, where two independent enzymes convert a common imaging substrate into spectrally distinguishable light. The signal intensity of one color is dependent upon biological state, whereas the other color is constitutively expressed. The ratio of emitted colored light corrects the functional signal for independent procedural variables, substantially improving the robustness and interpretation of relatively low-fold changes in functional signal intensity after drug treatment.

The application of these readouts *in vitro* is highly advantageous, as peak cell response to therapy can now be readily visualized for single or combination treatments and not simply assessed at an arbitrary and destructive timepoint. Spectral imaging *in vivo* can be challenging, but we also present evidence to show that the reporters can work in this context as well. Collectively, the development and validation of these internally controlled reporters allow researchers to robustly and dynamically visualize tumor cell biology in response to treatment. Given the prevalence of bioluminescence imaging, this presents significant and much needed opportunities for preclinical therapeutic development.

## Introduction

Bioluminescence imaging (BLI) with firefly luciferase is a popular approach in preclinical oncology and is frequently used to dynamically measure relative tumor cell viability *in vitro* or *in vivo* [1-4]. Its non-destructive nature allows tumor cells to be repeatedly measured over time in the same tissue culture sample or individual subject (e.g. before and after drug treatment). BLI has a large dynamic range and when employed in this manner it is not uncommon for signal intensity to change by several orders of magnitude over the course of an experiment. As an optical approach, BLI is not considered truly quantitative, but background imaging noise is low and these large changes in signal intensity over time can be meaningfully interpreted to reflect relative change in viable tumor burden.

Numerous examples also exist in the literature that describe creative transgenic strategies to restrict luciferase expression to a defined biological state (e.g. [5-7]). In this way, levels of bioluminescence can be used as a proxy indicator of functional biology and is therefore a powerful research tool to non-destructively assess specific effects of experimental genetic or pharmacologic perturbations. Such functional bioluminescent measures can be significantly more challenging to interpret however, as the difference in signal intensity pre and post experimental perturbation is often only several fold. Moreover, light output can be affected by multiple other factors unrelated to the biological measure, especially after the administration of a therapeutic. Between images, administered cancer drugs may reduce viable cell number, alter cellular metabolism or modulate drug efflux pump expression (and consequently substrate bioavailability [8,9]). Tumor hypoxia, pH [10], temperature (*in vitro*) [11], or depth in tissue (*in vivo*) [12] also affect light output. Moreover, a gain of functional bioluminescent signal is preferable, as signal loss can occur for technical reasons unrelated to the biology of interest.

To best address these issues and achieve robust and non-destructive measures of biological state, we describe here the development of two new and internally-controlled bioluminescent indicators of functional tumor cell stress. Called GemLuc and OxiLuc, these dual-color indicators confer non-destructive and corrected bioluminescent readouts of the cell cycle and oxidative stress respectively. We employ spectrally distinguishable green and red luciferase enzymes in these vectors (Click Beetle Green (CBG99) and *Photinus pyralis* red emitting luciferase 9 (PRE9) [13,14]). The protein stability of one color changes according to biology, whereas the other color is constitutively expressed and always on, acting crucially as an internal control to standardize and better interpret changes in the intensity of the functional measure. Both bioluminescent enzymes utilize the same imaging substrate (D-Luciferin), allowing correction of the functional signal for any variation in substrate bioavailability between images. Expression of the functional color is regulated at the post-translational level in both vectors and so they confer a fast response to experimental perturbation. A gain or loss of functional color relative to the constitutive expression of the internal control is now also of equal experimental relevance.

We present data here showing the application and validation of both internally-controlled readouts to robustly and non-destructively measure cycling cells or the induction of oxidative stress, as well as providing methodology for their correct use. To maximize ease of use, we have built both colored enzymes into a single lentiviral vector to deliver the stable expression of either imaging readout via a single cell transduction event. Our approach is versatile and readily customizable to other internally-controlled readouts of functional biology or bioluminescent enzyme.

## Results

### GemLuc and OxiLuc vector construction

To facilitate the introduction of the internally-controlled cell cycle (GemLuc) or oxidative stress (OxiLuc) readouts into a cell line of interest, each was built into a single lentiviral vector. As shown in figure 1a (and supplemental figure 1), both green luciferase (CBG99) and red luciferase (PRE9) expression cassettes are constitutively transcribed from two independent promoters.

**Figure 1.**
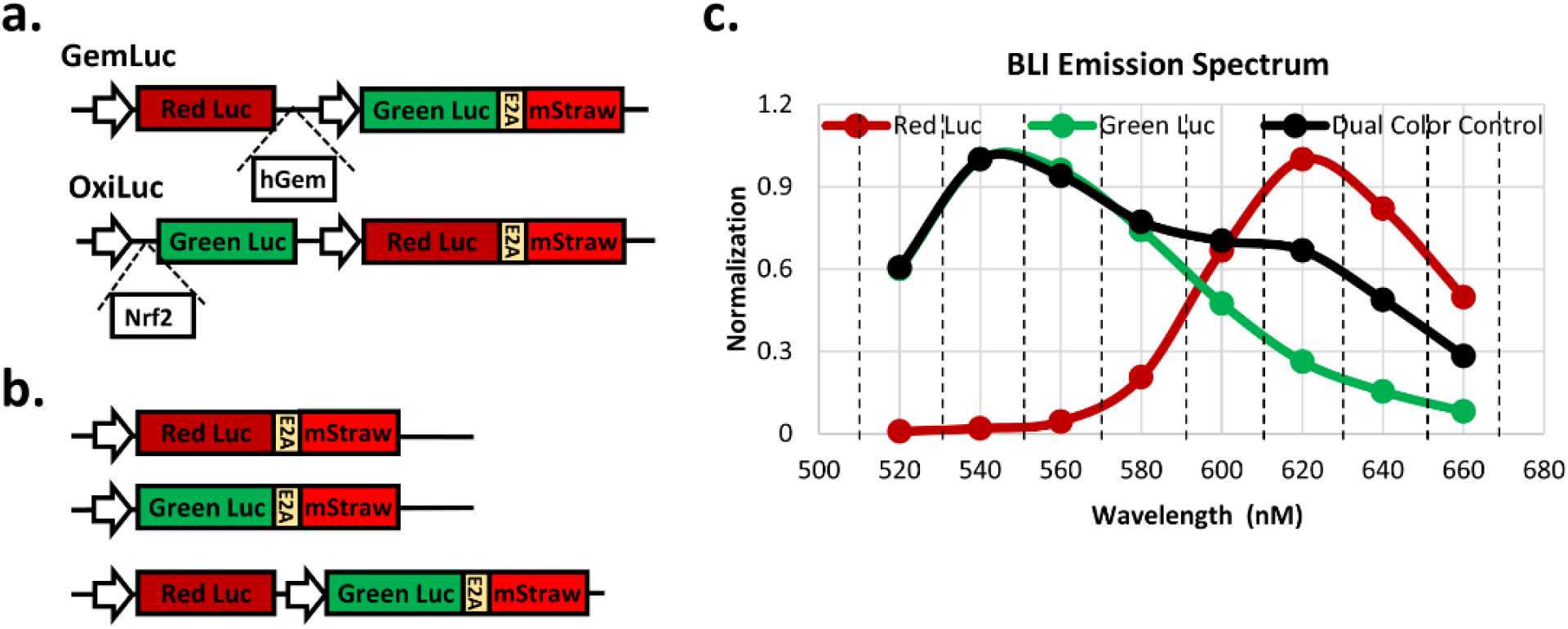
**a**. shows the transgenic organization of the internally-controlled cell cycle readout (GemLuc) and oxidative stress readout (OxiLuc). **b**. shows the transgenic organization of both the single-color control and dual-color control vectors. **c**. graph depicts spectral emission of green and red only control vectors *in vitro* (normalized to their peak emissions at 540 and 620 nm respectively), as well as the dual color control vector for reference. The dashed lines show the 20 nm bandwidth filters employed to acquire spectral emission.

To make GemLuc report cycling cells (cell cycle), we fused the Geminin degron (amino acids 1-110 of human Geminin protein) to the C-terminus of the red luciferase. This is the same peptide degron sequence employed in the original fluorescent FUCCI vector [15] and confers protein stability (and red bioluminescence) only at proliferative phases (S, G2 and M) of the cell cycle. The red luciferase is otherwise ubiquitinated and degraded by the proteasome at the non-proliferative G1 phase of the cell cycle. Relative to the “always on” green enzyme however, the presence or absence of red bioluminescence is meaningful, so the construct enables internally-controlled measures of both proliferative and non-proliferative phases of the cell cycle.

The green enzyme (CBG99) is approximately 3-fold brighter than the red enzyme (PRE9) and in an effort to boost OxiLuc imaging sensitivity, we reversed these colors such that the green enzyme is the functional color and red the internal control. We fused the NRF2 degron (Neh 2,4,5 domains) [16,17] to the N-terminus of the green enzyme. This conferred continuous proteasomal degradation of the green enzyme in “normal” cell conditions in a Keap1 dependent manner, but protein stabilization and an increase in green light relative to the red internal control in the presence of oxidative stress (OS). In both GemLuc and OxiLuc constructs, we positioned an E2A-mStrawberry cassette in-frame with the constitutively expressed internal control enzyme. This added the ability to sort stably transduced cells for red fluorescence by flow cytometry or to directly visualize transduced cells by fluorescence microscopy.

### Single color controls for spectral unmixing

There is considerable spectral overlap between the bioluminescent emissions of CBG99 and PRE9 (figure 1c). Accordingly, we used optical filters and a spectral unmixing algorithm [18] to best discriminate functional from control signal. To generate green or red only reference spectra and enable unmixing, we built constitutively-expressed single color lentiviral vectors (figure 1b). Both of these constitutively expressed either a PRE9-2A-mStrawberry or CBG99-2A-mStrawberry cassette from a constitutive promoter (CBH; short-form CAGGS) [19]. Polyclonal stably-transduced populations of MIA PaCa-2 cells for each color were then generated by flow sorting for strong fluorescence. Reference *in vitro* red and green bioluminescent spectra were then acquired by imaging these cells across a range of 20nm bandpass optical filters with an IVIS Spectrum scanner (Perkin Elmer). To demonstrate that our spectral unmixing approach was effective with these single-color reference spectra, we plated out mixtures of single red and single green expressing cells at defined ratios. Encouragingly, the fold ratio of measured and spectrally-unmixed red to green bioluminescence reflected the mixture of plated colored cells (supplemental figure 2).

### Non-destructive and internally-controlled readouts of treatment effects on the cell cycle (GemLuc)

To demonstrate the ability of GemLuc to indicate both proliferative and non-proliferative phases of the cell cycle, we added escalating doses of cisplatin (predicted S/G2 phase arrest [20]) or ribociclib (predicted G1/S phase arrest [21]) to a polyclonal population of GemLuc expressing MIA PaCa-2 cells *in vitro* and spectrally imaged 48hrs later. The results are shown in figure 2 and demonstrate that relative to expression of the internal control, GemLuc is able to non-destructively indicate the accumulation of cells in proliferative or non-proliferative phases of the cell cycle. Note that both chemotherapeutic treatments are partially cytotoxic and cause a decrease in signal intensity of the internal control (green light) in a dose dependent manner. By taking the ratio of red to green light however, we use the internal control to account for decreased viable cell number and correct the functional red signal, making the cell cycle measure more accurate. Flow cytometry of treated cells in repeated experiments validated the optical readout by showing comparable shifts of cells within different phases of the cell cycle after treatment (shown in supplemental figure3).

**Figure 2.**
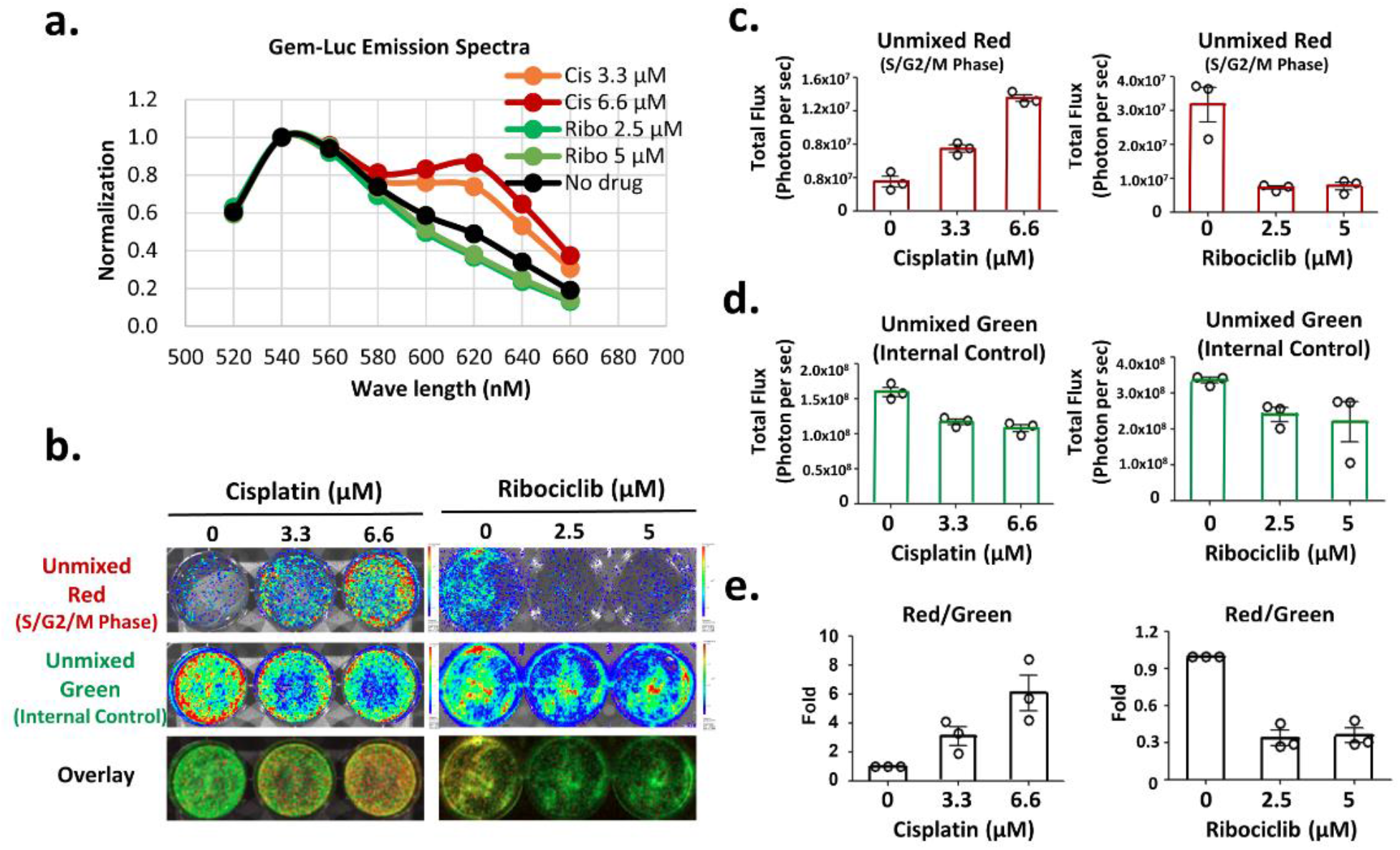
Non-destructive and internally-controlled readouts of treatment effects on the cell cycle (GemLuc). **a**. shows the normalized spectral emission of GemLuc expressing MIA PaCa-2 cells growing *in vitro* on different plates, 48 hours after drug treatment with either cisplatin or ribociclib. Spectra normalized to peak emission at 540 nm. **b**. representative spectrally-unmixed red and green images (and pseudocolor overlay) of GemLuc expressing MIA PaCa-2 cells growing *in vitro*, 48 hours after drug treatment. **c. d**. show measurement of functional red and internal-control green light after spectral unmixing and show both dose-dependent drug effects to the cell cycle and viable cell number. **e**. the ratio of measured red/green light corrects for experimental variables and produces a robust measure of effects to the cell cycle. Results show mean +/-SD and each individual data point of three technical replicates in one experiment.

### Visualizing cell cycle dynamics and synchronization after treatment

Our internally-controlled functional biology readouts are non-destructive and thus allow the researcher to follow dynamic effects of experimental perturbation over an extended period of time. We illustrate this in figure 3, where we treated GemLuc labelled MIA PaCa-2 cells *in vitro* with the CDK1 inhibitor RO3306 [22]. 24 hours after treatment and an evident cell cycle arrest at G2/M, one well was refreshed with normal media. These “released” cells are subsequently seen to progress synchronously into G1 phase of the cell cycle. Without an internal control to this measure, the loss of functional signal could have been attributed to cytotoxic effects of the drug, but normalization of the red functional signal with the green internal control corrects for this and we robustly see that surviving cells are synchronously entering a non-proliferative phase of the cell cycle. Maintenance of drug over this same time period was seen to perpetuate the G2/M arrest and give rise to a significant drop in cell viability at later experimental time points.

**Figure 3.**
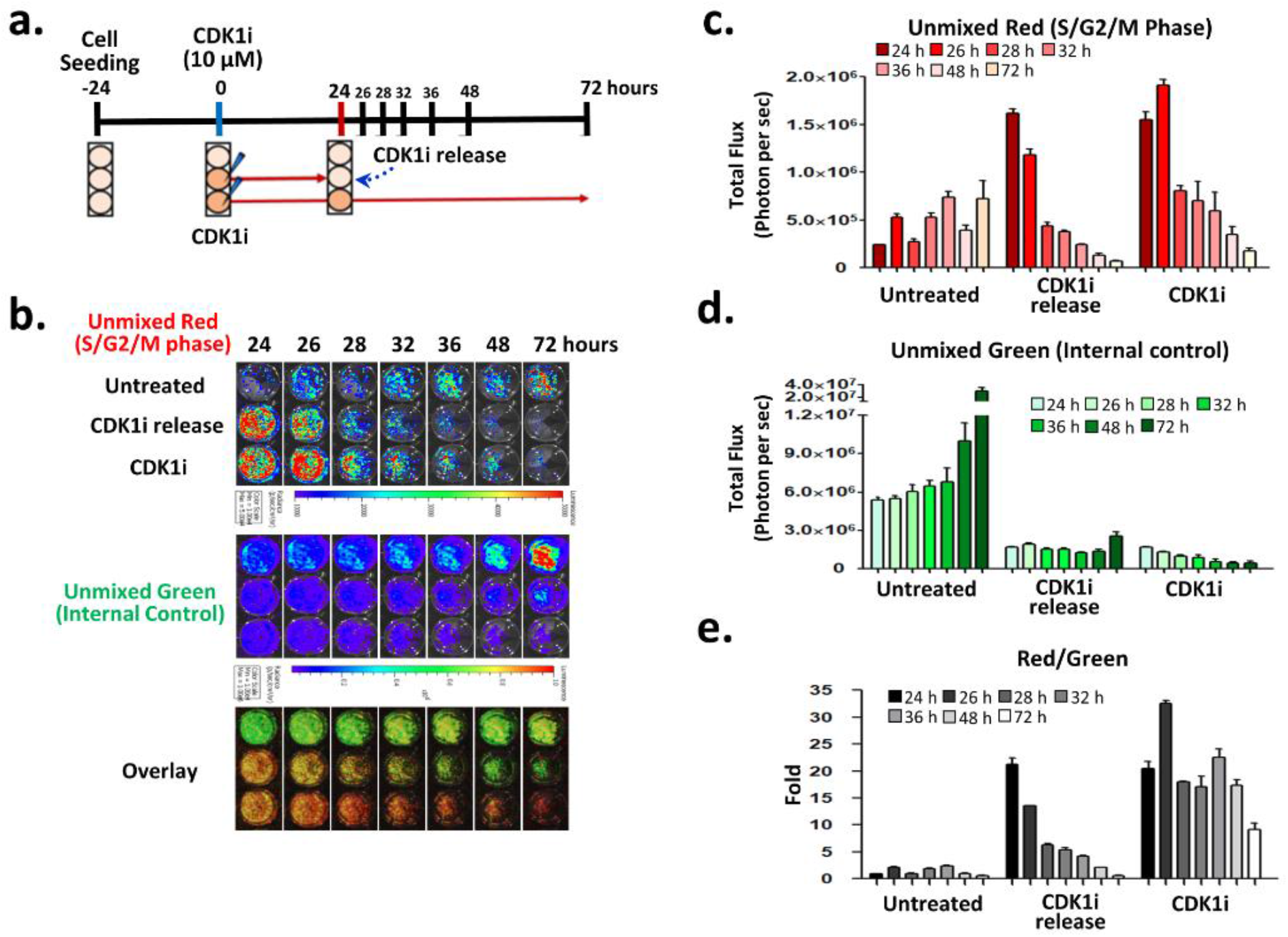
Imaging cell cycle dynamics. **a**. shows a schematic overview of the experiment involving the treatment of MIA PaCa-2/GemLuc cells with the CDK1 inhibitor, RO3306. **b**. shows spectrally unmixed or pseudocolored images of cells in the three experimental conditions, control untreated, 10 µM RO3306 treated for 24 hrs, then refreshed in regular tissue culture media, or 10 µM RO3306 treated for 72 hours. **c**., **d**. show spectrally unmixed measurements of functional red and internal-control green light, taken at regular timepoints over 72 hrs. **e**. the ratio of measured red/green light corrects the functional cell cycle readout and differentiates the cause of signal loss between differences in the proportion of cells in cycle versus viable cell number after treatment. Results show mean +/-SD of three technical replicates in one experiment.

### Non-destructive and internally-controlled readouts of treatment-induced OS (OxiLuc)

In parallel with the GemLuc experiments, we also tested the performance of the OxiLuc reporter for its ability to report physiologically-relevant levels of oxidative stress (i.e. intracellular ROS levels that stabilize Nrf2). The colors are reversed in this vector and induction of OS is indicated by an increase in green light. As shown in figure 4, we treated MIA PaCa 2-OxiLuc cells with escalating doses of Vitamin K3 (menadione), which is well known to induce ROS [23]. The fast response of our degron-based imaging readout was really exemplified in this study as we were able to measure a dramatic fold induction of the functional green color just 2 hours after administration of drug. The intensity of the control color was only a little affected as drug concentration increased over this short time period, resulting in minimal correction of the functional signal. Significantly however, we could substantially reduce both the induction of green light and the cytotoxicity of higher doses by repeating the experiment in the presence of the antioxidant NAC (N-acetylcysteine), indicating that the experimental observations are predominantly ROS dependent. We further confirmed ROS dependency by quantifying levels of HMOX-1 by qRT-PCR and the GSH/GSSG ratio on treated cell lysates (see supplemental figure 4). Similarly, we observed a robust induction of OS following the treatment of MIA PaCa 2-OxiLuc-cells after treatment with the investigational drug napabucasin (BBI-608) [24] (supplemental figure 5).

**Figure 4.**
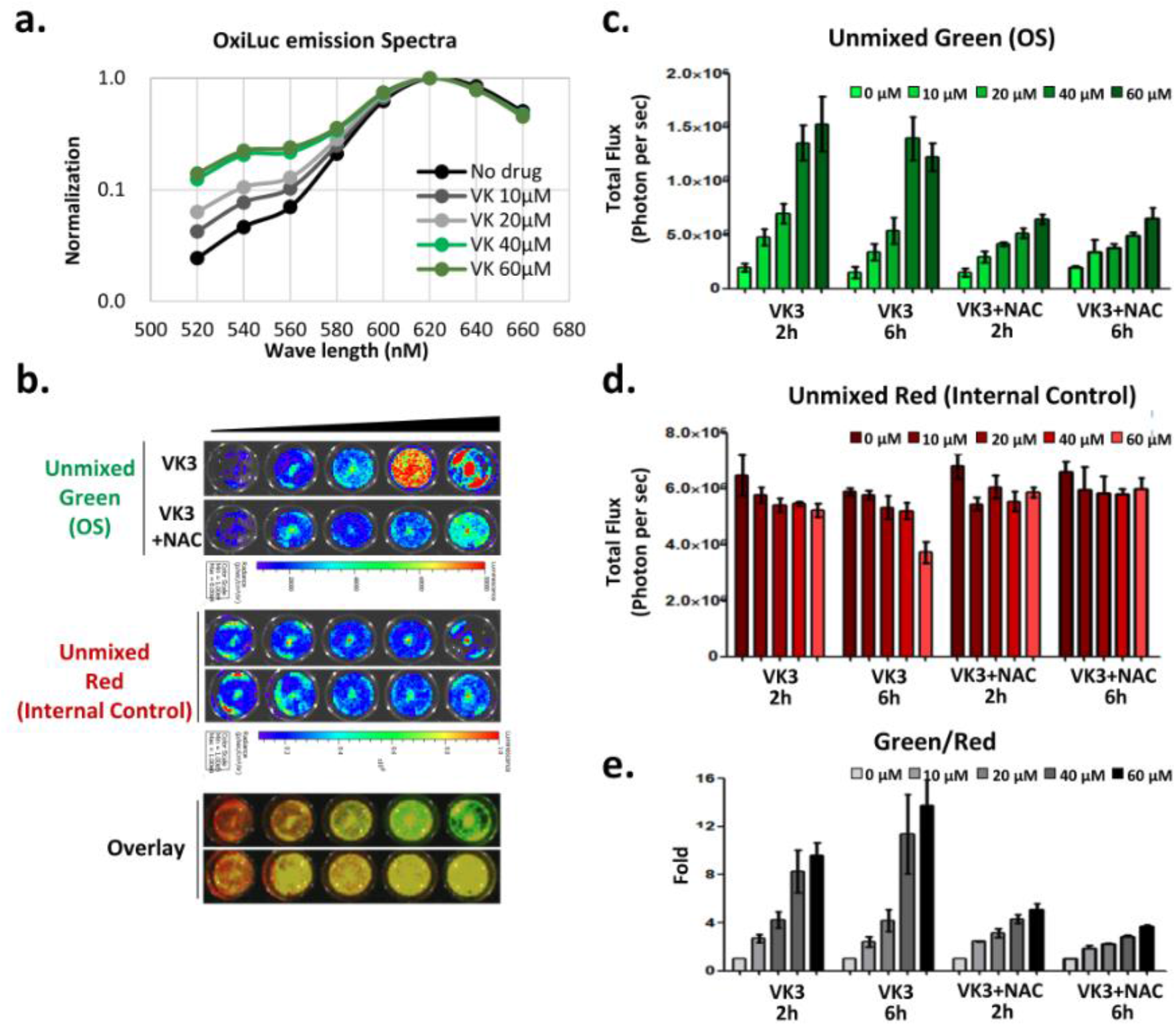
Non-destructive and internally-controlled readouts of treatment effects on the generation of OS (OxiLuc). **a**. shows normalized spectral emissions of OxiLuc expressing MIA PaCa-2 cells growing *in vitro* on the same plate, 6 hours after drug treatment with increasing concentrations of vitamin K3 (menadione). Spectra normalized to peak emission at 620 nm. **b**. representative spectrally unmixed red and green images (and pseudocolor overlay) of OxiLuc expressing MIA PaCa-2 cells growing *in vitro*, 6 hours after treatment with Vit K3 or Vit K3 and NAC. **c**.,**d**. show measurement of functional green (OS) and internal-control red light after spectral unmixing and show dose-dependent drug effects on the generation of OS (as early as 2 hours after treatment) and minimal effects on cell viability. **e**. the ratio of measured green/red light corrects for experimental variables and produces a robust measure of treatment effects on the induction of oxidative stress. Results show mean +/-SD of three technical replicates in one experiment.

Our *in vitro* imaging approach is amenable to relatively high throughput analysis and so can be useful in investigating the biological effects conferred by combination treatments. As shown in figure 5, we plated out OxiLuc expressing cells in a 6 by 6 grid on a 96 well plate format and varied the concentration of auranofin (Aur) along the y axis, and buthionine sulphoximine (BSO) along the x-axis. We observed minimal induction of OS with either drug alone. However, image analysis showed that the addition of even relatively low doses of BSO can potentiate the induction of oxidative stress in combination with doses of Aur above 100 nM.

**Figure 5.**
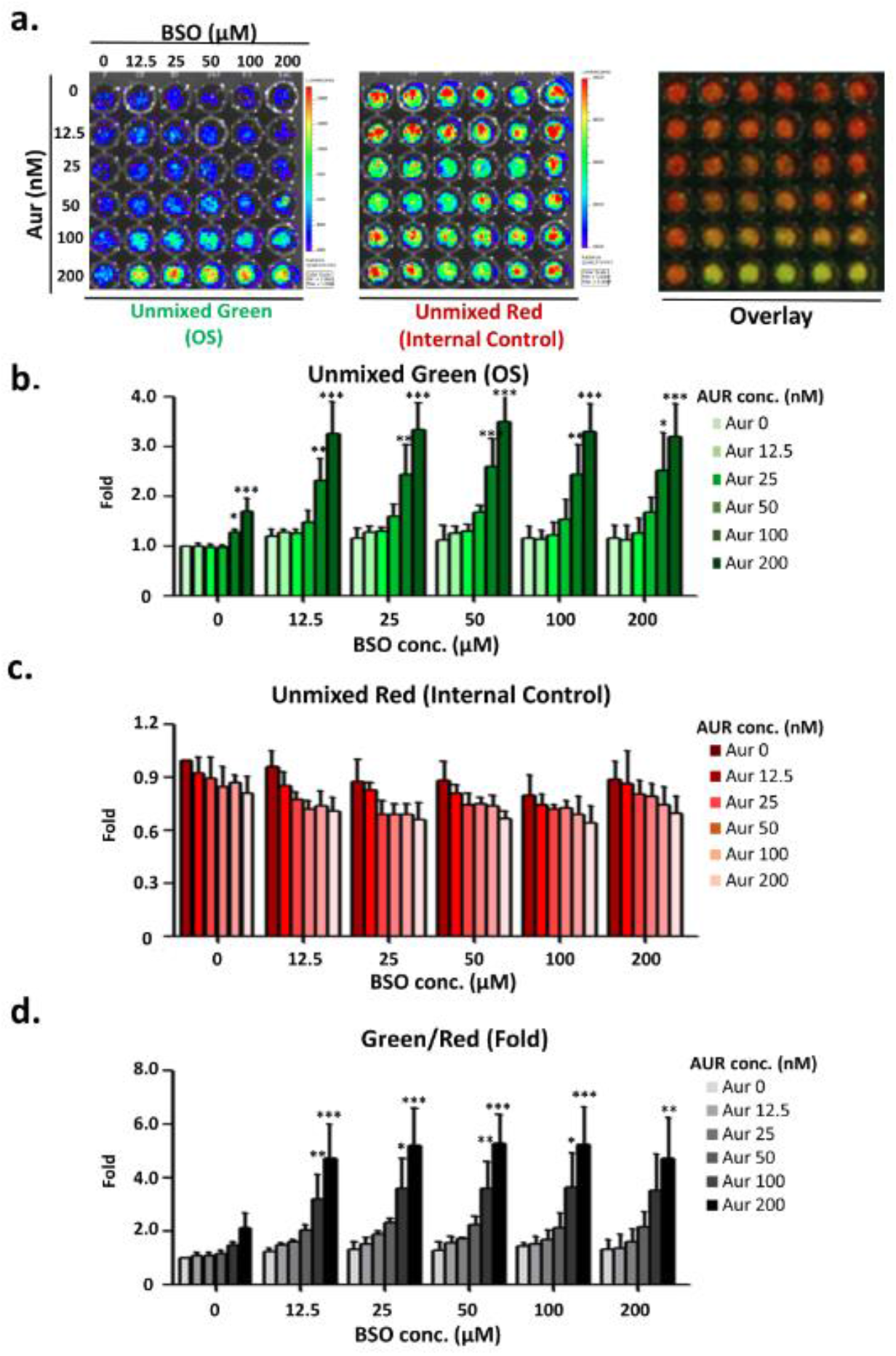
Imaging synergistic drug effects on the induction of OS. **a**. Spectrally-unmixed red and green images (and pseudocolor overlay) of OxiLuc expressing MIA PaCa-2 cells *in vitro*, plated out on a 96-well format, 24 hours after treatment with increasing concentrations of auranofin (top to bottom of plate) and buthionine sulphoximine (left to right of plate). **b**. Spectrally-unmixed green light was quantified following combination drug treatments (increased green light indicative of OS). **c**. Quantified measures of spectrally-unmixed red light (internal control) taken in parallel to correct the functional measure for experimental variables, resulting in **d**., a green/red fold ratio and robust non-destructive measure of OS induction. Results show mean +/-SD of three independent experiments. The significance of differences between the groups were compared to the baseline for each BSO concentration using ANOVA, followed by pairwise comparisons using Welch’s two-sample t-test. P-values were adjusted for multiple comparisons by Dunnett’s method.

### *In vivo* imaging of functional biology

BLI is particularly well-suited to making sensitive non-invasive *in vivo* measurements. However, spectral BLI *in vivo* can be challenging and is complicated by the fact that green light is significantly less transmissible through tissue than red light [25]. We therefore reasoned that labeled tumors should develop at both a superficial and fixed body location to maximize *in vivo* performance. Accordingly, we established single-color reference spectra from subcutaneous tumors to allow spectral-unmixing in future experiments (see supplemental figure 6). We then conducted two key control experiments to show that the internal-control can broadly correct the functional color for two important *in vivo* experimental variables. To demonstrate correction of the functional measure for differences in viable tumor cell number between images, we set up 6 mice with subcutaneous GemLuc expressing MIA PaCa-2 tumors and spectrally imaged them at regular time intervals over an 18-day period (supplemental figure 7). Without an internal control, the intensity of the functional color increased more than 40-fold as the tumors grew in size and cellularity increased, but it is not possible to attribute the contribution an increase in the proportion of cells in S, G2 and M have made to this increase. The functional cell cycle measure could largely be corrected by the internal control color however, suggesting relative stability in the proliferation rate over this extended period of tumor growth.

To show that the internal control could also correct the functional signal for variable substrate bioavailability *in vivo*, we took repeated spectral measures of GemLuc expressing tumors following a single I.P. injection of substrate. It is well established that FLuc bioluminescence rises rapidly to peak levels between 10 and 15 minutes after I.P. injection of D-Luciferin substrate, then slowly decreases again to baseline over the course of the next 2 hours. Those changes in bioluminescent intensity over time are primarily dependent on substrate bioavailability. As shown in supplemental figure 8, we essentially observe those dynamic changes to functional color intensity, but again, those are not related to the proportion of proliferating cells. Once corrected by the internal control color, we see relatively steady state light output from the functional color over this extended period of time, irrespective of substrate levels.

To demonstrate the ability of OxiLuc to image the induction of OS *in vivo* following the administration of a drug (see figure 6), we implanted 6 mice with subcutaneous OxiLuc expressing MIA PaCa-2 tumors. Once tumors had visibly developed, we separated all mice into two cohorts; one treated with sodium arsenite (ASN) to induce OS [26] and the other vehicle control. Based on *in vitro* observations, mice were spectrally imaged before and 5 hours after treatment.

**Figure 6.**
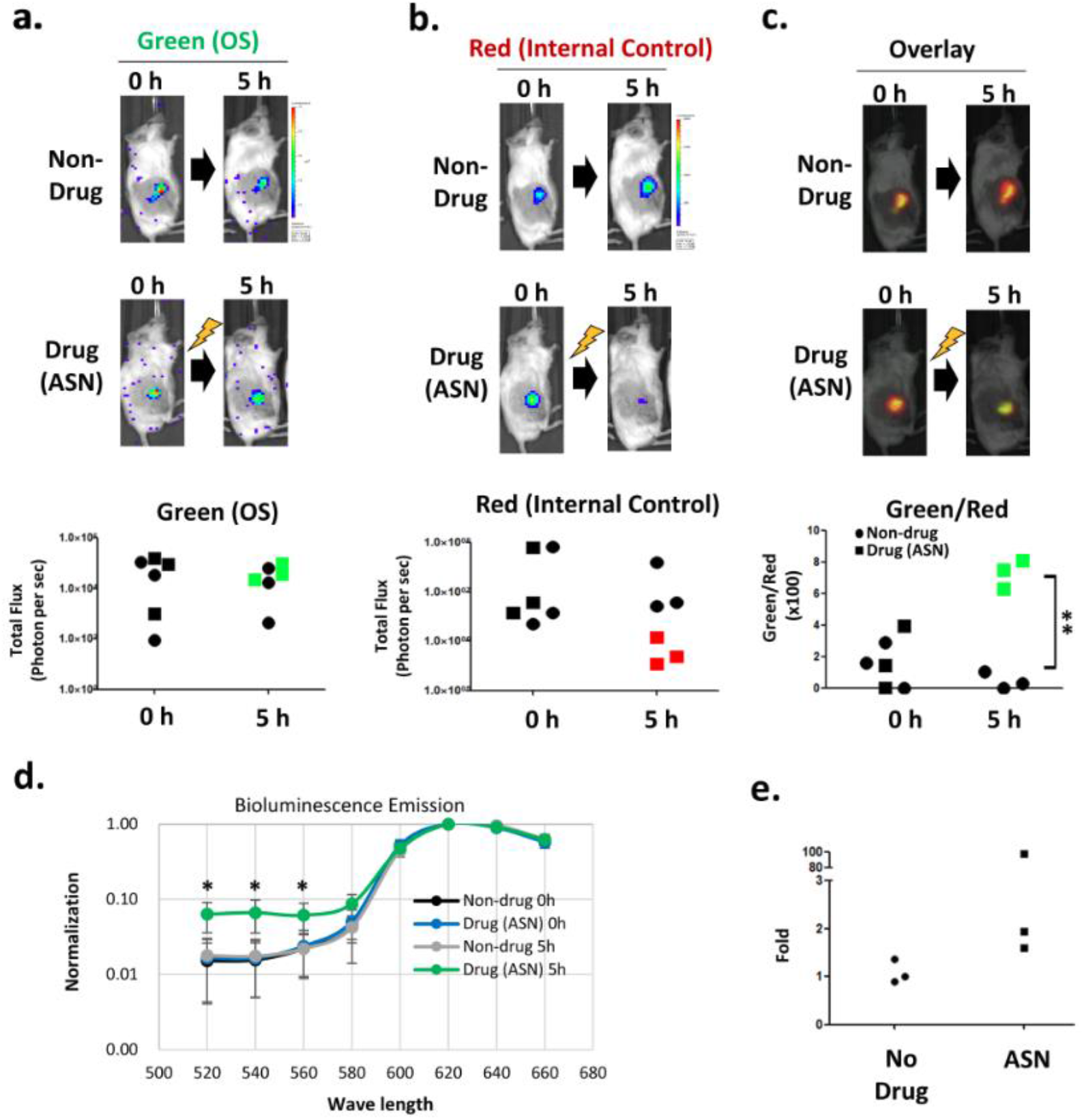
Imaging the induction of oxidative stress *in vivo*. Subcutaneously engrafted OxiLuc expressing MIA PaCa-2 tumors were treated with either sodium arsenite (n=3) or saline control (n=3). **a**. A representative image of spectrally-unmixed green light (indicative of OS) before and 5 hours after treatment. **b**. Spectrally-unmixed red light (internal control) at the same timepoints and **c**. A pseudo-color overlay showing that ASN treated tumors have a significantly higher fold-increase in the ratio of green to red light than the saline treated group at this 5-hour timepoint (** p<0.01). A two-sided Student’s t-test was used to identify the difference between groups. **d**. Spectral emission (normalized to 620 nm) from both experimental cohorts, before and after treatment. Two-way ANOVA was used to identify the difference between groups. (* p<0.05). **e**. The results of qRT-PCR analysis on HMOX expression to validate the induction of OS in treated mice at this single 5-hour timepoint.

The results of this experiment were encouraging; measurement of the functional green color alone showed little evidence of an increase following treatment with OS-inducing ASN relative to untreated tumors. However, light emission from the internal control red enzyme was markedly decreased in treated mice and so when the ratio of green to red light was made, ASN treated mice showed an induction of OS, which was validated by analysis of HMOX 1 RNA expression levels by qRT-PCR.

## Discussion

Bioluminescence is frequently employed in preclinical research and drug development as a means to non-destructively measure dynamic aspects of functional tumor biology. However, given the number of non-specific variables that can substantially influence light output, it can be challenging to confidently interpret low-fold changes to bioluminescence in response to treatment, especially *in vivo*. Cancer therapeutics are intended to exert tumor cell specific stress and death, which in turn also affect cellular metabolism and the tumor microenvironment, all of which can have unpredictable effects on signal output unrelated to the biological measure. This was dramatically illustrated in two classic papers that showed that certain cancer treatments affect the expression levels of the ABCG2 drug efflux pump [8], (or P-glycoprotein for coelenterazine metabolizing enzymes [27]), which in turn significantly affected intracellular substrate levels and gave rise to surprising shifts in bioluminescent intensity post-treatment.

In an effort to greatly improve the robustness of functional bioluminescent measures, which unarguably have an important role to play in preclinical research and drug development, we report here the construction of two dual-color and internally-controlled readouts of functional biology. Named GemLuc and OxiLuc (for detection of changes in the cell cycle or in levels of oxidative stress respectively), the functional signal is corrected from substantial experimental variability by a constitutively expressed control color that metabolizes the same substrate. Others have previously described the application of two or more spectrally distinct bioluminescent colors to simultaneously image multiple independent biological parameters in mouse models [14,28,29]. Although technically impressive, none of the multi-parametric measures are internally controlled and so the signal from each one is potentially prone to influence from any of the various factors that influence light production.

Unlike the well-defined and spectrally tight light emission typical of fluorophores, the emission wavelengths of CBG99 and PRE9 are broad and overlap considerably. Consequently, we generated single-color reference spectra to facilitate spectral unmixing and best attribute emitted photons to either colored enzyme. Both GemLuc and OxiLuc vectors are relatively modular however and it should be straightforward to employ different colored enzymes with alternative imaging characteristics, or different regulatory elements for alternative functional biology readouts.

The *in vitro* application and dynamic nature of these readouts is straightforward and highly advantageous experimentally. Without imaging, it is common for researchers to choose a single or small number of arbitrary timepoints after treatment (e.g. at 24 hours) to assess drug effects. These new internally-controlled readouts of proliferation or oxidative stress now confer robust and temporal pharmacodynamic measures, amenable to moderately high-throughput (96 well format), allowing the researcher to better visualize both how and when a cell responds to single or combination treatment prior to recovering or succumbing to cell death. As stability of the functional color is regulated at the post-translational level in both vectors, we can observe fast responses to treatment (e.g oxidative stress within 2 hours).

*In vivo* imaging of colored light is complicated by the differential tissue transmission properties of red and green light. However, we determined that tumors developing at a fixed and relatively superficial body location could be effectively imaged in this way. Following *in vivo* demonstration that our approach broadly corrects the functional signal for two potentially major variables between scans (differences in viable cell number and substrate bioavailability), we showed the importance of an internal control to appropriately interpret images of OS induction following drug treatment of OxiLuc expressing tumors. Sodium arsenite is well-known for its ability to induce ROS *in vivo*, however, quantitation of the functional color alone, 5 hours after treatment, showed no meaningful difference in emitted light intensity between treated and untreated tumors. Quantitation of the internal control color however showed a significant decrease in light output in treated mice and the corrected green/red light ratio could clearly differentiate the induction of OS in treated mice. We have not exhaustively tested the *in vivo* sensitivity of this approach, as that is highly context dependent and subject to the model, reporter transgene expression levels and tumor-size. However, we note that *in vivo* imaging sensitivity may be enhanced in the future with different bioluminescent enzymes with different spectral light emission properties. With these vectors and current imaging hardware, we believe that the utility of our approach will be limited to imaging relatively superficial tumors *in vivo*.

It should also be noted that the measured fold change of the functional signal relative to control is not an absolute measure and is relative to the intensity of baseline signal. For example, we have observed that in comparison to the MIA PaCa-2 cells described here, OxiLuc labelled SUIT-2 pancreatic cancer cells possess higher constitutive baseline levels of oxidative stress (data not shown). As a result, despite showing similarly high levels of functional bioluminescence upon the induction of oxidative stress, they exhibit a comparably lower fold induction. Similarly, you would expect a highly proliferative cell line to have more intense baseline functional color than a slow growing line and so in absolute terms, a lower fold induction after a drug-induced S/G2/M cell cycle arrest. The ability to observe robust and dynamic changes to tumor cell biology following treatment is not compromised though.

In conclusion, the validated internally-controlled reporters described here confer robust bioluminescent measures of anti-cancer drug effect beyond routine measures of cell viability. Their non-destructive nature mean that peak levels of growth arrest or oxidative stress induction can now be observed dynamically and not measured at an estimated arbitrary and destructive timepoint.

## Materials and Methods

### GemLuc and OxiLuc plasmid construction

all gene expression constructs were built into the pBOB lentiviral backbone (3^rd^ generation lentiviral vector [30]) to facilitate subsequent labeling and stable expression of our reporters in eukaryotic cell lines of interest. Reporter genes (CBG99, PRE9 and mStrawberry), protein degron sequences, additional genetic elements (e.g E2A sequence [31]) and useful flanking restriction sites were all synthesized from GenScript USA (New Jersey) and assembled as finished vectors (as shown in figure 1 and supplemental figure 1) within 1 to 3 restriction enzyme or Gibson Assembly cloning steps. All restriction enzymes, T4 DNA ligase, taq DNA polymerase and Gibson cloning kit were supplied by NEB. All cloning and sequencing primers were supplied by Sigma (Burlington, MA). All plasmids were grown up in Top10 competent bacteria (Invitrogen, Waltham, MA) and plasmid DNA prepped from bacteria with Qiagen DNA mini prep or maxi prep kits Germantown, MD).

### Single color control plasmids

single color red or green lentiviral vectors were built to generate reference spectrum for subsequent spectral unmixing. As shown in figure 1b, lentiviral vectors either contained CBG99-E2A-mStrawberry or PRE9-E2A-mStrawberry expression cassettes, transcribed from a constitutive CBH promoter (short-form CAGS promoter). Stably transfected cells were positively selected for mStrawberry fluorescence by FACS (BD LSR Fortessa™ Cell Analyzer, BD Bioscience, Franklin Lakes, NJ) or fluorescence microscopy.

### *In vitro* cell culture, lentiviral packaging and reporter labelling

HEK293T cells (ATCC, Manassas, VA) and MIA PaCa-2 cells (human pancreatic cancer; ATCC, Manassas, VA) were maintained in Dulbecco’s modified Eagle’s medium (Corning, Lowell, MA) supplemented with 10% fetal bovine serum (Corning, Lowell, MA) and Penicillin-Streptomycin (Corning, Lowell, MA). All cells were grown at 37°C and 5% CO2 in a humidified incubator and tested negative for mycoplasma using the MycoAlert Mycoplasma Detection Kit (LT07-318, Lonza).

Lentivirus was produced by transfection of HEK293T packaging cells with three packaging plasmids (pMDL, pRSV-REV, pCMV-VSVG) [30] and the lentiviral reporter plasmid using Lipofectamine 3000 (Invitrogen, Waltham, MA) according to the manufacturer’s recommended protocol. Supernatant containing lentivirus was then collected 72 hours later, aliquoted and stored at -80°C.

MIA PaCa-2 cells were seeded on a 6 well plate at a density of 3×10^5^ cells/well and transduced with 500 µl lentiviral supernatant plus polybrene to a final concentration of 8 μg/ml. Transduced cells were then expanded further in culture and FACS sorted for mStrawberry expression (BD-FACS Aria, BD Bioscience, Franklin Lakes, NJ).

### Spectral bioluminescent imaging and drug treatment *in vitro*

all bioluminescence imaging was conducted with an IVIS Spectrum scanner (Perkin Elmer, Shelton, CT). Spectral bioluminescent images of live cells growing *in vitro* were acquired 2 minutes after adding 150 ug/ml of D-Luciferin (Goldbio, Saint Louis, MO). Sequential images were acquired through optical filters ranging from 520 to 660nm at 20nm intervals, using the Spectral Unmixing option in the Imaging Wizard of Living Imaging software 4.0 (Perkin Elmer, Shelton, CT). For Gemluc and cell cycle imaging, exposure times were 10 sec for each filter. For OxiLuc and oxidative stress imaging, to compensate for low levels of green light, 60 second images were acquired from 520 to 580 nm optical filters, then 30 sec from 600 to 660 nm. All *in vitro* images were acquired at field of view C and focused at 0.5 cm subject height. Photon flux from each well was quantified as photons/second/steradian/cm^2^ with Living Image Software 4.0 (Perkin Elmer, Shelton, CT). A spectral unmixing algorithm within Living Image, trained with our single-color reference spectra, was then applied to better discriminate the bioluminescent light from CBG99 (green) or PRE9 (red) enzymes.

For cell cycle imaging, 3×10^5^ MIA PaCa-2/GemLuc cells were either plated day -1 on a black-walled 6 well plate (Cellvis, Mountain View, CA) or a clear bottom 6 well plate (Corning, Glendale, AZ).

Bioluminescent spectra were then acquired from day 0, before and after treatment with cisplatin or ribociclib (Selleckchem, Radnor, PA), as indicated in the main text. For dynamic imaging of the cell cycle, 5×10^4^ MIA PaCa-2/GemLuc cells were seeded on a black-walled 24 well plate (Cellvis, Mountain View, CA) and treated with 10 µM of RO-3306 (CDK1 inhibitor; Sigma, Burlington, MA) for 24 hours. The culture media was then refreshed and imaged as indicated in the main text.

For oxidative stress imaging, 5×10^4^ or 8×10^3^ MIA PaCa-2/OxiLuc cells were plated day -1 on a black-walled 24 well plate or 96 well plate (Cellvis, Mountain View, CA). Bioluminescent spectra was then acquired after VK3 and NAC (N-acetylcysteine) treatment or auranofin and buthionine sulfoximine (both Sigma, Burlington, MA) treatment or BBI-608 (Boston Biomedical, Cambridge, MA) and Dicoumarol (Selleckchem, Radnor, PA) treatment, as indicated in the main text.

### Flow cytometry

to quantifiably analyze the cell cycle readout, treated cells were harvested at defined experimental time points and fixed with 2% formaldehyde in PBS for 15 min at room temperature. Fixed cells were then washed with PBS and stained with 3 µM DAPI (Thermo Scientific, Waltham, MA). Cells were then analyzed with a FACSAria flow cytometer (BD Bioscience, Franklin Lakes, NJ) to determine the proportion of cells in G1 or S, G2, M phases of the cell cycle. Cellular DNA content was measured with a UV trigon detector array with a 450/50 band-pass filter and data analyzed by Flow Jo software (BD Bioscience, Franklin Lakes, NJ). At least 10,000 events were collected per sample.

### Quantitative RT-PCR

total RNA was extracted from tumor cells (before or after drug treatment) with TRIzol reagent (Invitrogen, Waltham, MA). cDNA was then synthesized using 1 mg of total RNA and M-MLV Reverse Transcriptase according to manufacturer’s instructions (Invitrogen, Waltham, MA).

Quantitative PCR was then performed at 95°C for an initial 3 min followed by 40 cycles of denaturation at 95°C for 15s, annealing and extension at 60°C for 60s using *Hmox-1* specific Taqman primer (Applied Biosystems, San Francisco, CA) and TaqMan Universal Master Mix (Applied Biosystems, San Francisco, CA) on a QuantStudio 6-flex Real time-PCR instrument (Applied Biosystems, San Francisco, CA). Quantification of relative gene expression was performed with the QuantStudio Real-Time PCR software v1.1 (Applied Biosystems, San Francisco, CA) and normalized to actin expression levels.

### Quantification of glutathione

The GSH/GSSG ratios were quantified using the GSH/GSSG-Glo assay kit (Promega, Madison, WI) to measure intra cellular oxidative stress. To measure total glutathione or GSSG of the cell, each sample was tested in 96-well culture plates with total glutathione or GSSG lysis reagent. After the 5 minutes lysis reaction, the luciferin generation reagent and detection reagent were added. The plates were then read in a SpectraMax Microplate reader (Molecular devices, San Jose, CA).

### *In vivo* tumor models and *in vivo* spectral bioluminescent imaging

all procedures were conducted in accordance with the IACUC at Cold Spring Harbor Laboratory and all mice were housed in a temperature and humidity-controlled vivarium on a 12 hr dark-light cycle with free access to food and water.

All *in vivo* studies employed either male NSG (NOD.Cg *Prkdc*^*scid*^*Il2rg*^*tm1Wji*^/SzJ) or nude mice (Balb/c *Foxn1*^*nu*^*/Foxn1*^*nu*^), aged between 6–8 weeks old (JAX laboratory, Bar Harbor, ME). All *in vivo* bioluminescence imaging was conducted with an IVIS Spectrum scanner (Perkin Elmer, Shelton, CT) To establish tumors, 1×10^6^ MIA PaCa-2 cells (expressing GemLuc or OxiLuc (or single color controls to train spectral unmixing)), were engrafted subcutaneously. Once tumors had visibly developed (≥5 mm in diameter), mice were spectrally imaged and treated with cancer therapies as indicated in the main text. Briefly, mice were anesthetized with 2% isoflurane and injected I.P. with 300 mg/kg of D-Luciferin (Goldbio, Saint Louis, MO). Anesthesia was maintained and spectral images were acquired starting at 15 minutes after injection. As per *in vitro* spectral imaging, sequential images were acquired through optical filters ranging from 520 to 660nm at 20nm intervals, using the Spectral Unmixing option in the Imaging Wizard of Living Imaging software 4.0 (Perkin Elmer, Shelton, CT). Exposure time per filter was 10 sec or 60 sec for cell cycle imaging or oxidative stress imaging respectively, using field of view D and focused at 1.5 cm subject height. Living image software 4.0 (Perkin Elmer) was then used to analyze and spectrally-unmix images when appropriate. The signal intensity of emitted light was quantified as photons/second/steradian/cm^2^ by Living Image Software 4.0 (Perkin Elmer, Shelton, CT). A spectral unmixing algorithm within Living Image, trained with our single-color reference spectra, was then applied to better discriminate the bioluminescent light from CBG99 (green) or PRE9 (red) enzymes.

To induce of oxidative stress *in vivo*, 10 mg/kg of sodium arsenite (ASN; Sigma, Burlington, MA) or PBS control was injected intraperitoneally. All mice were spectrally imaged immediately before and 5 hours after ASN treatment.

## Statistical analysis

Statistical analyses were performed using R Statistical Software (version 4.0.4; R Foundation for Statistical Computing, Vienna, Austria), Graphpad Prism 5 (GraphPad software, Inc, La Jolla, CA), and Microsoft Excel (Redmond, WA). The p-value less than 0.05, 0.01 and 0.001, mentioned as *, **, *** respectively, are considered as statistically significant.

## Acknowledgements

The authors would like to thank Pam Moody and the Flow Cytometry shared resource at CSHL for technical assistance with all flow cytometry and cell sorting. We would also like to thank the Animal shared resource at CSHL for assistance with all aspects of animal husbandry and Catherine Winchester (CRUK Beatson Institute, Glasgow, UK) and Taehoon Ha (biostatistician, CSHL) for guidance on data presentation and statistical analysis.

