## Supplemental Figures for "Internally-controlled and dynamic optical measures of functional tumor biology"

**Supplemental figure 1.**

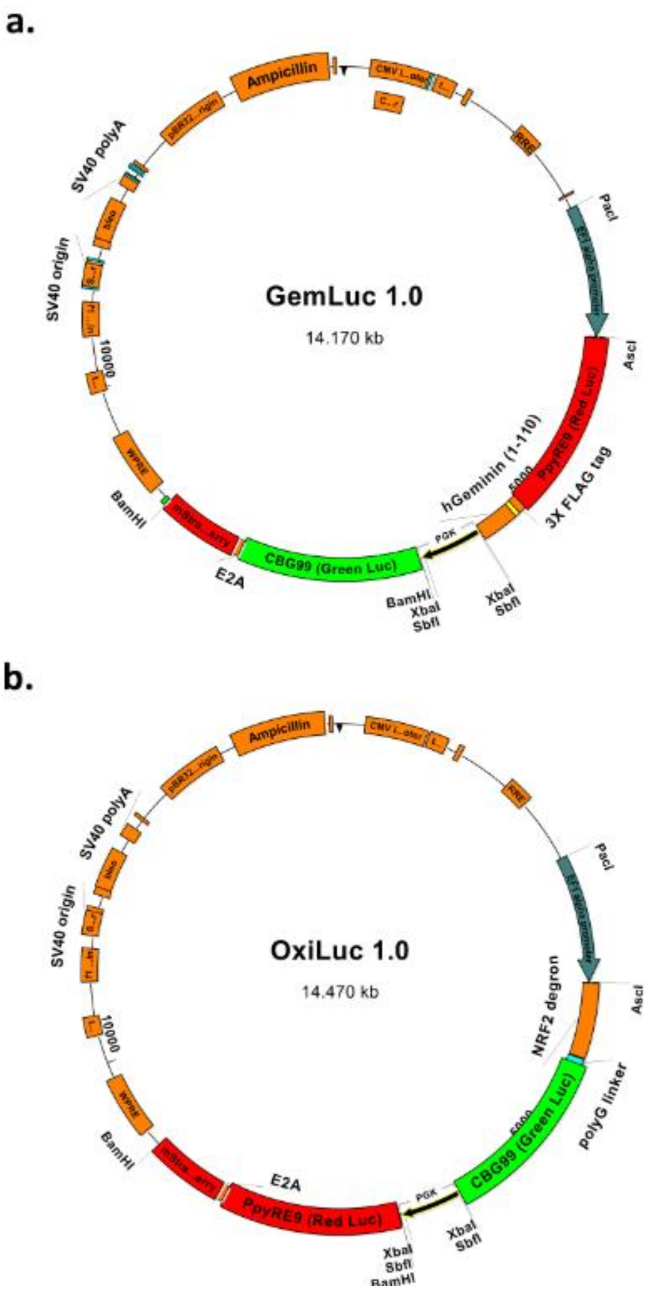


**Supplemental figure 1.**

Detailed plasmid maps of (**a**) GemLuc and (**b**) OxiLuc lentiviral vectors with important restriction enzyme sites marked.

**Supplemental figure 2.**

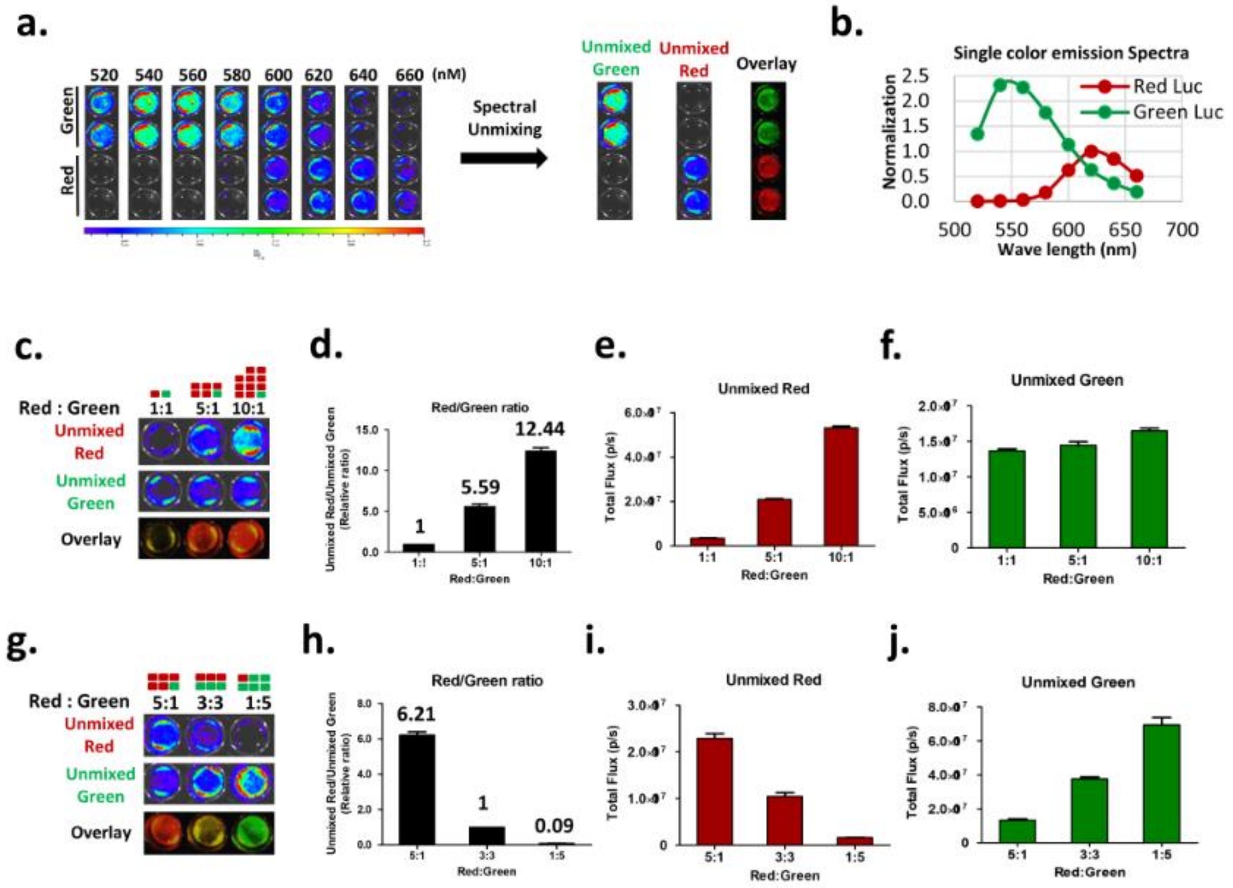


**Supplemental figure 2**

**a.** Multispectral image acquisition of MIA PaCa-2 cells expressing single color red or green luciferase control vectors. These were then used as reference spectra for spectral-unmixing in other experiments. **b**. Quantitation of single-color emission spectra. **c.-f.** and **g.-j.** Defined mixtures of single color red or green expressing MIA PaCa-2 cells spectrally imaged *in vitro*. The ratio of measured red to green light broadly recapitulates the mixture of plated cells. Results show mean +/- SD of three technical replicates in one experiment.

**Supplemental figure 3.**

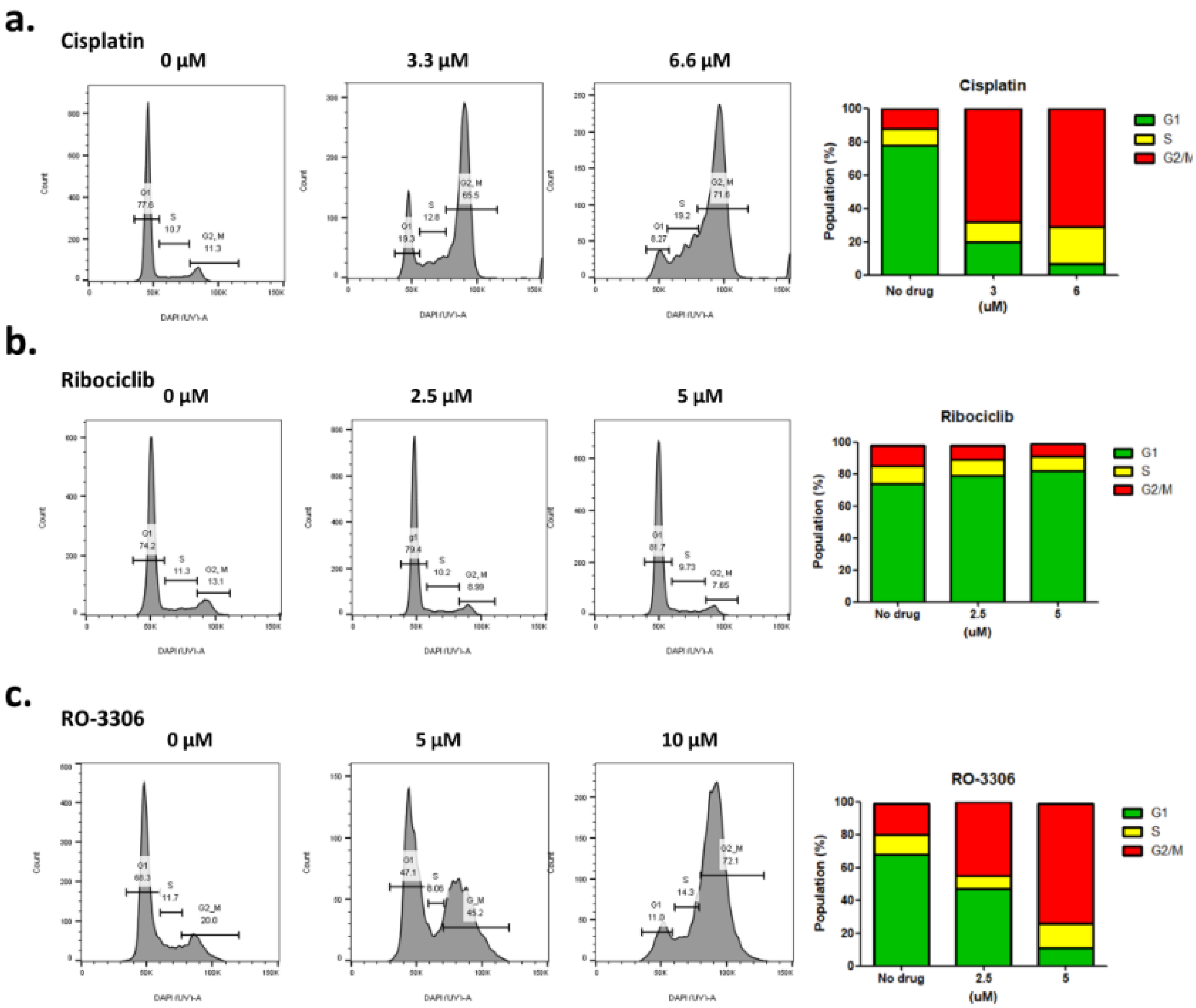


**Supplemental figure 3.**

Quantification by flow cytometry of the percentage of GemLuc expressing MIA PaCa-2 cells in G1, S, or G2/M phases of the cell cycle after *in vitro* treatment with (**a.**) cisplatin, (**b.**) ribociclib or (**c.**) RO-3306, validating the optical imaging readout.

**Supplemental figure 4.**

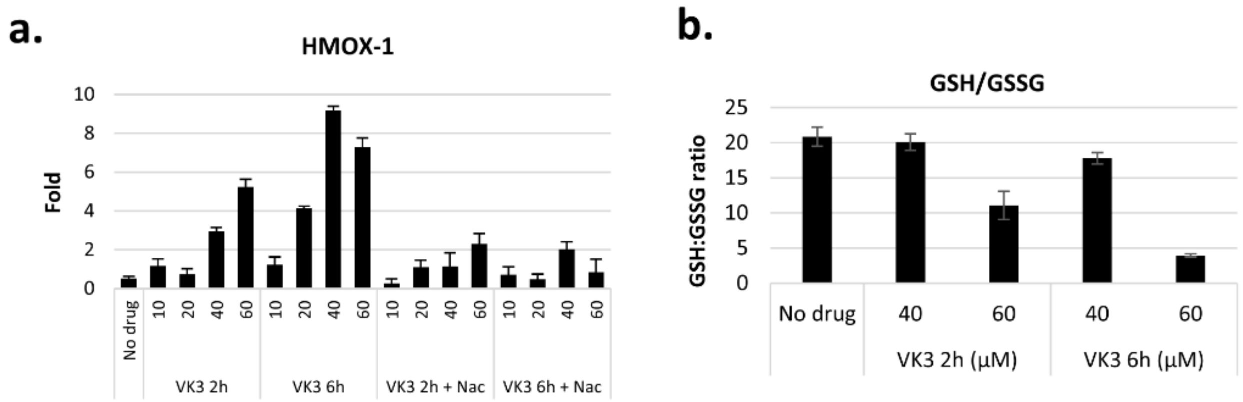


**Supplemental figure 4.**

**a.** Validation of ROS induction in OxiLuc expressing MIA PaCa-2 cells by qRT-PCR analysis of Heme Oxygenase 1 (HMOX1) expression levels following vitamin K3 treatment *in vitro*. **b.** ROS induction also confirmed by a decrease in the ratio of reduced glutathione (GSH) to oxidized glutathione (GSSG). Results show mean +/- SD of three technical replicates in one experiment.

**Supplemental figure 5.**

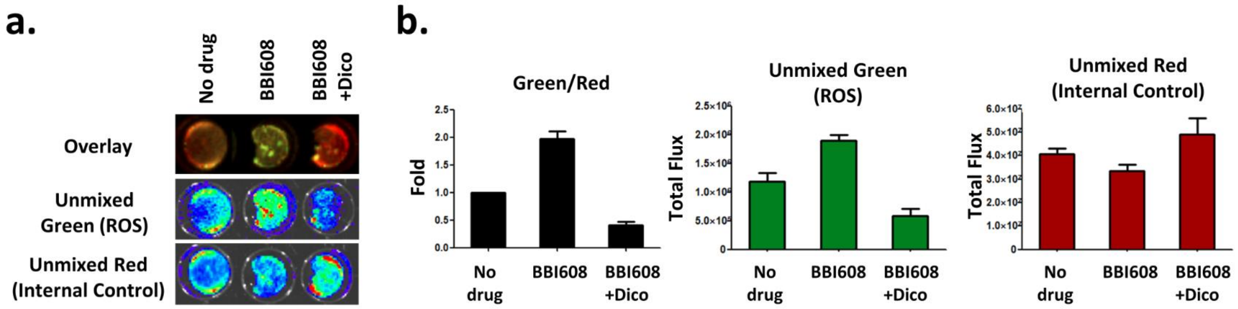


**Supplemental figure 5.**

Imaging napabucasin (BBI-608) drug effects on the induction of OS. **a.** Spectrally-unmixed red and green images (and pseudocolor overlay) of OxiLuc expressing MIA PaCa-2 cells *in vitro*, plated out on a 24-well format, 24 hours after treatment with 0.5 µM of BBI-608 as a single agent, or in combination with anti-oxidant dicoumarol (Dico) at 10 µM. **b**. Quantitation of spectrally-unmixed green and red light after drug treatment; increased green light relative to red indicates OS induction. Results show mean +/- SD of three technical replicates in one experiment.

**Supplemental figure 6.**

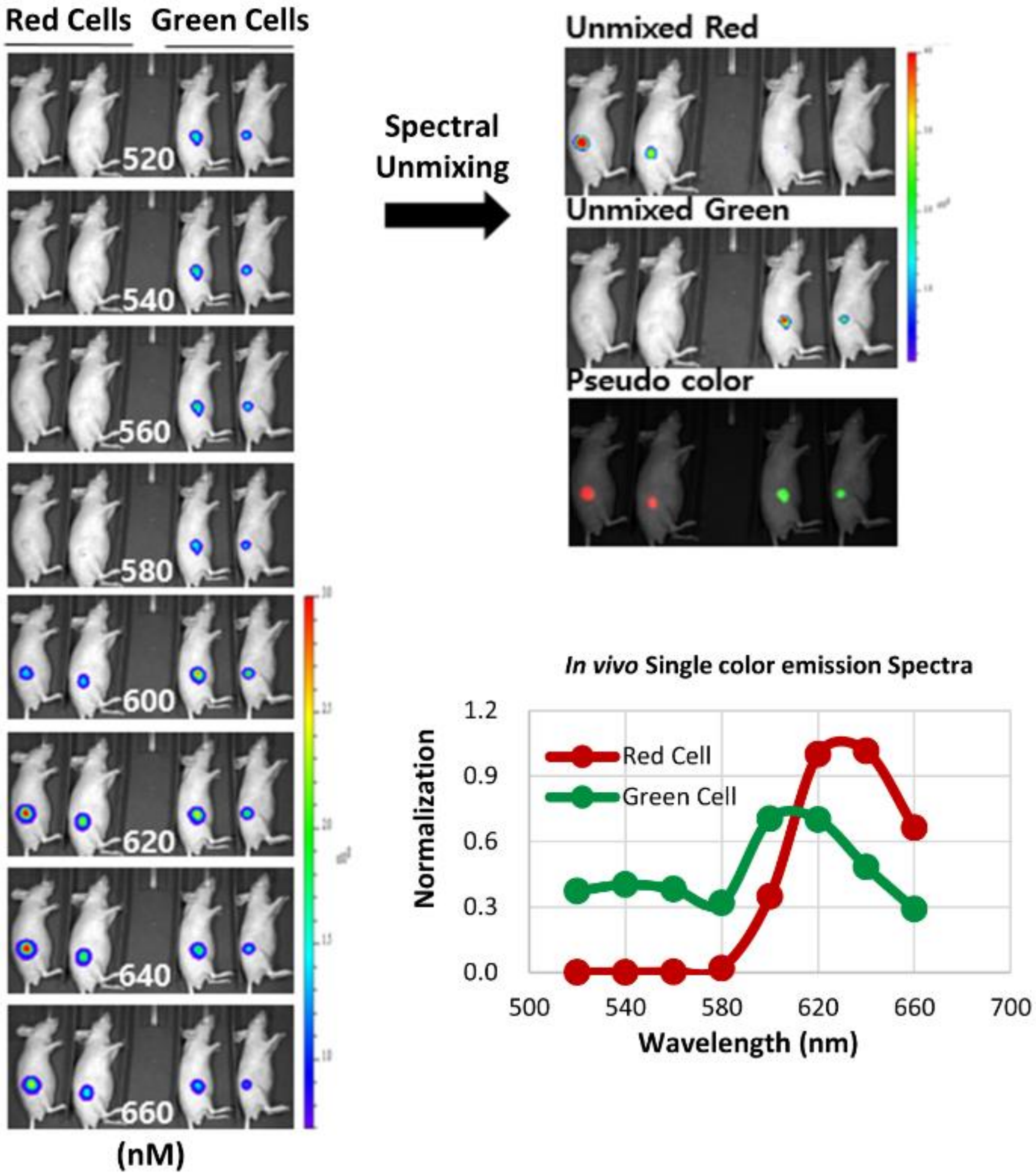


**Supplemental figure 6.** Shows the acquisition of reference red and green spectra from single-color expressing MIA PaCa-2 cells developing as subcutaneous tumors *in vivo*; to enable spectral-unmixing in subsequent dual-color experiments.

**Supplemental figure 7**

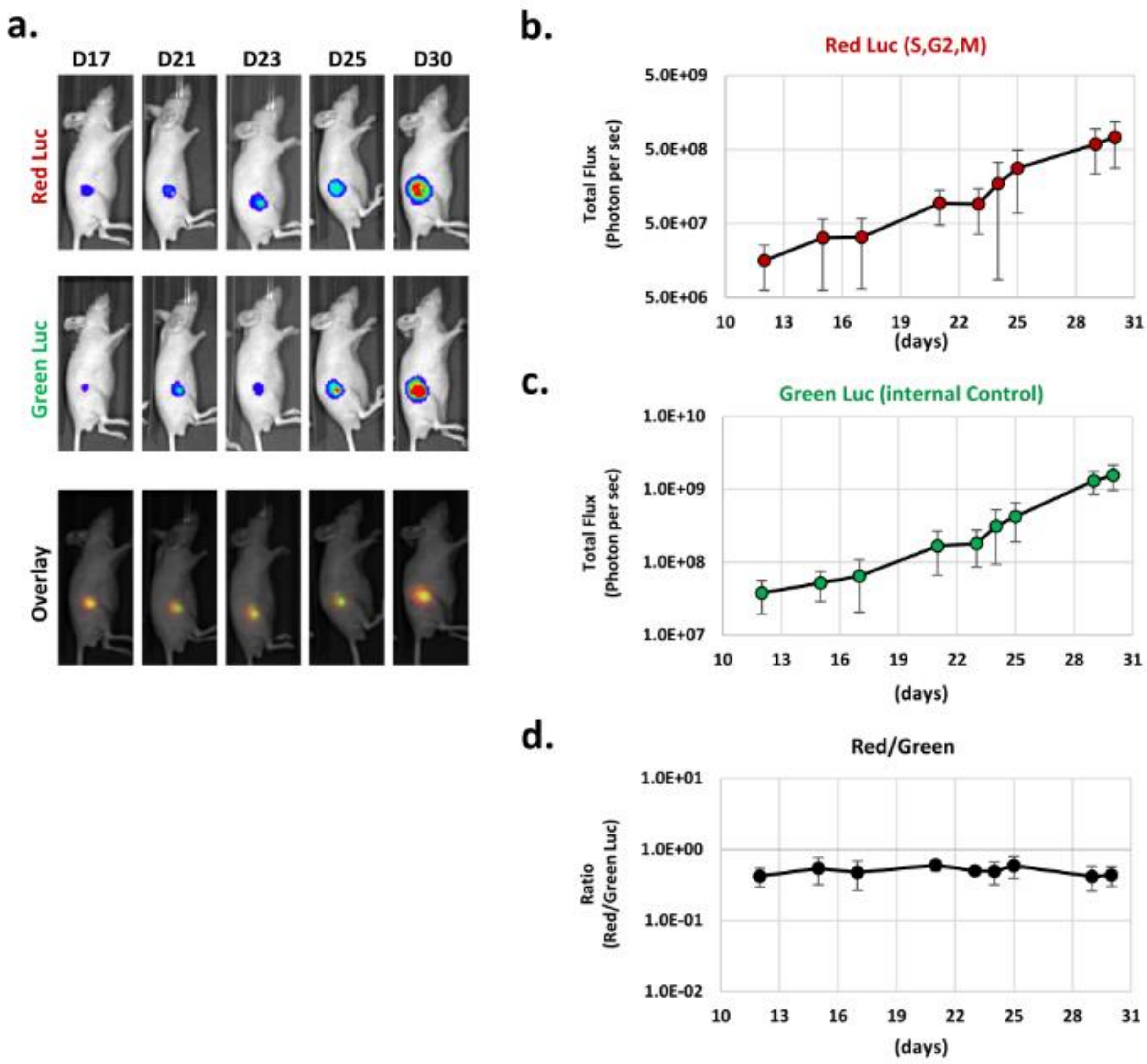


**Supplemental figure 7.** Figure showing that the measured ratio of red/green light *in vivo* largely corrects for changes in viable tumor cell number over an extended period of 18 days and >40 fold difference in the intensity of functional color emission over this same time period

**Supplemental figure 8**

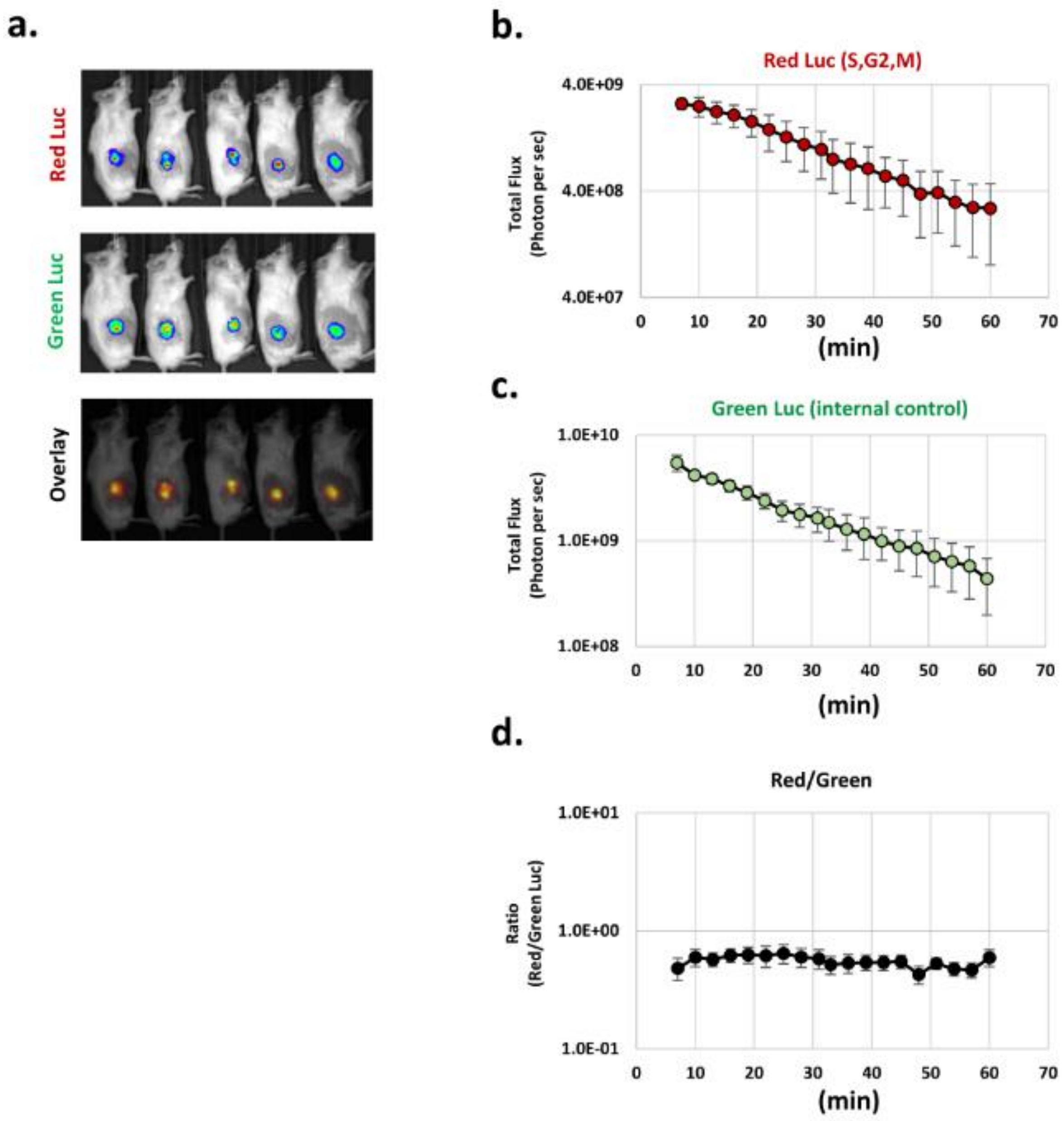


**Supplemental figure 8.** Figure showing that the measured ratio of red/green light *in vivo* largely corrects for broad changes in the level of bioavailable imaging substrate, D-Luciferin, despite photon/second levels of the single functional color changing approximately 10-fold over the same time period.

### Supplementary table 1.
### Proportion of emitted light from Red Luc or Green Luc in each filters

| | 520 | **540** | 560 | 580 | 600 | **620** | 640 | 660 |
|---|---|---|---|---|---|---|---|---|
| **Red Luc** | 0.25% | **0.60%** | 1.36% | 6.31% | 20.45% | **30.64%** | 25.15% | 15.24% |
| **Green Luc** | 14.04% | **23.38%** | 22.47% | 17.36% | 11.09% | **6.15%** | 3.61% | 1.90% |

### Supplementary table 2.
### Proportion of emitted light from Gem-Luc in each filters after drug treatmen

| | 520 | **540** | 560 | 580 | 600 | **620** | 640 | 660 |
|---|---|---|---|---|---|---|---|---|
| **No drug** | 12.4% | **20.5%** | 19.3% | 15.1% | 12.0% | **10.0%** | 6.9% | 3.9% |
| **Cis 3.3 uM** | 11.1% | **17.7%** | 16.3% | 13.6% | 13.4% | **13.1%** | 9.4% | 5.4% |
| **Cis 6.6 uM** | 9.8% | **16.5%** | 15.7% | 13.4% | 13.7% | **14.2%** | 10.6% | 6.2% |
| **Ribo 2.5 uM** | 14.1% | **22.4%** | 20.6% | 15.5% | 11.1% | **8.2%** | 5.3% | 2.9% |
| **Ribo 5 uM** | 13.0% | **22.0%** | 20.9% | 16.0% | 11.3% | **8.3%** | 5.5% | 3.0% |

### Supplementary table 3.
### Proportion of emitted light from OxiLuc in each filters after drug treatment

| | 520 | **540** | 560 | 580 | 600 | **620** | 640 | 660 |
|---|---|---|---|---|---|---|---|---|
| No drug | 0.73% | **1.40%** | 2.11% | 6.35% | 18.74% | **30.07%** | 25.43% | 15.18% |
| VK 10uM | 1.24% | **2.25%** | 3.01% | 7.30% | 19.72% | **29.29%** | 23.31% | 13.88% |
| VK 20uM | 1.79% | **2.98%** | 3.64% | 7.99% | 19.86% | **28.22%** | 22.12% | 13.40% |
| VK 40uM | 3.22% | **5.31%** | 5.59% | 8.83% | 18.89% | **25.80%** | 20.27% | 12.10% |
| VK 60uM | 3.55% | **5.66%** | 6.05% | 9.04% | 18.81% | **25.33%** | 20.05% | 11.49% |

**Supplemental tables.** **1.** Spectral quantitation of emitted light from single-color control expressing cells, (as shown in supplemental figure 2). **2**. Spectral quantitation of emitted light from GemLuc expressing cells following drug treatment (as shown in figure 2). **3**. Spectral quantitation of emitted light from OxiLuc expressing cells following drug treatment (as shown in figure 4).
